# Regionally specific picture naming benefits of focal tDCS are dependent on baseline performance in older adults

**DOI:** 10.1101/2024.11.25.625288

**Authors:** A. Yucel, M. Meinzer, A. K. Martin

## Abstract

Word-finding difficulty is a common challenge in older age and is linked to various neuropathological conditions associated with ageing. Transcranial direct current stimulation (tDCS) has shown promise as a cognitive enhancement tool for both healthy aging and age-related cognitive disorders. However, its effectiveness in enhancing word-finding ability remains inconsistent, especially among healthy adults. This variability is likely due to factors such as task selection, stimulation parameters, and small sample sizes. Additionally, many studies have overlooked the role of baseline performance in evaluating tDCS efficacy. In this preregistered study, we examined 72 older and 72 younger adults using a double-blind, sham-controlled design, delivering anodal focal tDCS to either the left inferior frontal gyrus or the left temporoparietal junction. Baseline naming performance and fluid intelligence were measured before stimulation. Anodal stimulation of the left inferior frontal gyrus significantly increased response speed for object and action naming in older adults, but crucially only in older adults who performed poorly during the baseline naming session. Findings demonstrate regionally specific effects of focal tDCS in healthy older individuals in greater need for naming facilitation. Notably, performance on a broad measure of fluid intelligence was unrelated to stimulation response, suggesting task specificity of this effect.

## Introduction

Difficulty finding words is a common complaint in advanced age (James & Burke, 2000; Wei et al., 2024) and is exacerbated in several neurodegenerative conditions (Cotelli et al., 2007; Ralph et al., 1998), with considerable impact on quality of life and economic burden (Falck et al., 2022). An ageing population in Western countries only heightens the importance of improving cognitive outcomes into advanced age with novel interventions the focus of considerable scientific endeavour (Sabayan et al., 2023). One such method that has been proposed, both in healthy ageing (Perceval et al., 2016) and neurodegenerative conditions (Cammisuli et al., 2022; Darkow et al., 2017; Roncero et al., 2019; Wang et al., 2023), is transcranial direct current stimulation (tDCS). TDCS involves the application of a weak electrical current over target brain regions to increase or decrease neural excitability, via modulation of the resting membrane potential (Stagg & Nitsche, 2011). As a clinical application and research tool, tDCS holds considerable promise due to the minimal costs involved, the portability of the device, and the relative ease of use (Brunoni et al., 2012).

Language function, including word retrieval, is often targeted for modulation using tDCS, although considerable heterogeneity of outcomes are reported in healthy cohorts (Westwood & Romani, 2017). There are several factors that likely lead to these inconsistent results, including the use of different target regions, delivery of stimulation either prior to (offline) or during (online) task performance, and the type of naming task used (e.g., lexical decision, verbal fluency, confrontation naming). Moreover, chronological age and associated changes in brain structure and function, as well as baseline performance levels have been shown to impact on tDCS effects (Meinzer et al., 2024; Perceval et al., 2016).

Therefore, the present study systematically investigated tDCS effects using a common word-retrieval task (i.e., picture naming) in young and older adults and stimulated two key hubs of the task-relevant brain network, the left inferior prefrontal gyrus (lIFG) and the left temporoparietal junction (lTPJ), either before or during the task to investigate offline and online effects of tDCS. Moreover, we use a focal tDCS method allowing region-specific neural modulation by constraining the current to the targeted regions (Gbadeyan et al., 2016; Martin, Huang, et al., 2017, 2019; Martin, Su, et al., 2019). Finally, we investigated potential modulating effects of baseline cognitive performance (i.e., naming performance and a measure of general fluid intelligence) on the degree of tDCS effects. This allowed investigating the task specificity of potential baseline performance effects on tDCS response.

Naming ability is considered a good proxy for language competence and although reliant on semantic, phonological, and articulatory processing, there is consistent evidence associating performance with activity across a broad left hemispheric fronto-temporo-parietal network (Indefrey & Levelt, 2004; Piai & Eikelboom, 2023). TDCS studies targeting the left inferior frontal gyris (IFG), have returned mixed results in healthy, predominantly young, cohorts (see Westwood & Romani, 2017). In older adults, evidence is scarce. A prominent study identified facilitation of picture naming in older adults after left inferior frontal stimulation coupled with decreased neural activity at the stimulation site, but in a sample consisting of only ten older adults without a control stimulation site (Holland et al., 2011). Another study used an offline, bilateral stimulation protocol to the IFG and claimed an order-specific effect in improving naming performance only when active stimulation was delivered in the second session (Lifshitz-Ben-Basat & Mashal, 2018). The small sample and exploratory order-dependent effects are not sufficient to conclude the effectiveness of tDCS to improve naming performance in healthy older adults. Likewise, further studies have identified a facilitation of proper name retrieval following left temporal stimulation (Ross et al., 2010) or left dorsolateral stimulation in older adults, but only for online stimulation (Fertonani et al., 2014). Again, both relied on small samples and only looked at stimulation effects in one region, limiting power and site-specific evidence. Therefore, claims that tDCS may be effective in improving picture naming in healthy adults and potentially preventing or treating neurodegenerative conditions associated with ageing (Siegert et al., 2021), requires more robust, well-powered evidence.

Despite inconsistent results of tDCS in healthy cohorts (Westwood & Romani, 2017), evidence is more consistent in cohorts with brain damage. For example, positive effects on naming ability have been shown in aphasic patients (Cappon et al., 2016; Crinion, 2016; Sandars et al., 2016), progressive supranuclear palsy (Madden et al., 2019), and dementia (Roncero et al., 2019). Although, even in clinical populations, reviews have called for more robust evidence (Al Harbi et al., 2017). More consistent stimulation effects in clinical cohorts, and to some extent healthy ageing, suggests that stimulation may be more effective on brains with suboptimal levels of cortical excitability within task-specific networks. Ageing-related cognitive decline is not uniform and it is possible that stimulation interventions may only be beneficial in those who demonstrate lower cognitive performance, indicative of suboptimal brain function relevant to the specific cognitive task. Baseline-dependent effects of stimulation have been identified in older adults in the domains of associative learning (Perceval et al., 2020), visual detection (Learmonth et al., 2015), and word generation (Meinzer et al., 2013), but as yet have not been investigated in relation to naming ability. A recent review highlighted baseline cognitive performance as the key factor in determining the effectiveness of tDCS in older adults, with stimulation showing the greatest positive effects in those with lower baseline cognitive function (Koo et al., 2023), motivating the inclusion of baseline functioning in the current study.

Although baseline cognitive functioning may be the key determinant, other factors may influence stimulation effects on naming ability in older adults. For example, stimulation can be delivered online during the administration of the task or offline, preceding task administration. Briefly, the neuronal mechanism underpinning online stimulation effects relate to the depolarization of the cell via the influx of sodium ions into the cell, causing an initial depolarization. This depolarization then activates voltage-gated calcium channels of the NMDA receptor. The changes in membrane potentials will affect the excitability of neurons in a manner that depends on the polarity of the stimulation. Offline tDCS effects are thought to reflect the early stages of the neuroplastic response, primarily mediated by the activation of NMDA receptors, which is triggered by a reduction in GABAergic tone. A reduction in GABA levels within the stimulated region beneath the electrode has been noted following anodal tDCS (Friehs & Frings, 2019; Stagg & Nitsche, 2011; Stagg et al., 2009). In the domain of naming, one previous study identified online specific effects in older adults (Fertonani et al., 2014), but in a small sample and with stimulation to the left dorsolateral prefrontal cortex. Differential effects of tDCS in young and older adults depending on timing of stimulation have also been shown, with young adults demonstrating immediate effects of stimulation, whereas older adults showed a delayed response of 20 minutes, thought to reflect age-related changes in neuroplasticity (Fujiyama et al., 2014). The current study will extend this evidence to stimulation effects at two key hubs of the left-lateralized naming network in a large cohort of healthy young and older adults and assess stimulation effects during both the immediate, online period and the delayed, offline period.

Naming tasks reliably activate a broad fronto-temporo-parietal network (Indefrey & Levelt, 2004), but targeting specific regions within the network, such as the lIFG and the lTPJ may influence the stimulation response. First, the two regions may serve subtly different processes relevant to naming. For example, action naming has been associated with greater frontal activity, compared with greater temporal activity associated with object naming (Vigliocco et al., 2011). Several studies using tDCS have supported a dissociation of stimulation response, with lIFG stimulation improving action naming and lTPJ stimulation improving object naming (Fiori et al., 2011, 2013; Marangolo et al., 2013). However, the parsing of nouns and verbs in English is especially problematic due to the fact the words often do not differ morphologically (*hammer* – could act both as verb or noun) and many verbs are often created from nouns (e.g. *the chair, to chair)*. Moreover, evidence suggests that neural activity dissociated on the basis of semantic association rather than grammatical class (Scott, 2006). Although specific targeting of regions associated with semantic association is beyond the scope of the current study, a failure to show dissociable effects of lIFG and lTPJ stimulation on action and object naming would provide tentative support for semantic association rather than grammatical class as the key dissociable neural process.

Therefore, in the present preregistered study https://osf.io/wsmxt) we aim to provide evidence for the efficacy of focal tDCS to the lIFG and lTPJ to improve object and action naming in healthy young and older adults. Moreover, we assess differences attributable to administering tDCS online or offline and look at the modulating influence of baseline naming and general cognitive performance on subsequent stimulation effects.

## Materials and Methods

### Participants

Participants were neurologically healthy, right-handed, native English speakers who were tDCS-naive. Young adults were aged 18–30 years, and older adults were aged 55–85 years. Both young and older adults were stratified to receive either lIFG or lTPJ stimulation in a sham-controlled, repeated-measures design. The young IFG group (25F/11M), young TPJ group (28F/8M), older IFG group (22F/14M), and older TPJ group (27F/9M) were comparable in terms of gender distribution, χ²(3, *N* = 171) = 2.82, *p* = 0.42.

The mean age of the older adults in the lIFG group (*M* = 71.8 years, *SD* = 6.1) and the lTPJ group (*M* = 71.4 years, *SD* = 7.2) was comparable, *t*(70) = 0.25, *p* = 0.81, *d* = 0.06. Among young adults, those who received lIFG stimulation (*M* = 19.6 years, *SD* = 2.2) were slightly older than those in the lTPJ group (*M* = 18.7 years, *SD* = 1.2), *t*(70) = 2.29, *p* = .02, *d* = 0.24.

The older groups were matched on education level (data were missing for three older adults): lIFG group (*M* = 15.9 years, *SD* = 2.0) and lTPJ group (*M* = 15.3 years, *SD* = 2.4), *t*(67) = 1.06, *p* = .30, *d* = 0.25. All young adults were current undergraduate students.

Older adults were recruited from the University of the Third Age (U3A) and the community surrounding The University of Kent. The older adults were provided with £15 as compensation for their time. Young adults were recruited from the University of Kent and the surrounding community and received either course credits or £15 compensation for their time.

Standard tDCS safety guidelines for exclusions were followed (Antal et al. 2017 Clin Neurophysiol), and included history of seizures or migraines, metallic objects in the head, electrical medical equipment on the person that was unable to be removed. None of the participants reported using medicines that are known to interfere with the effects of tDCS. The study was conducted in compliance with the ethical standards outlined in the Declaration of Helsinki. All procedures involving human participants were reviewed and approved by the University of Kent Psychology Research Ethics Committee [ID: 202216587800677857], and informed consent was obtained from all participants prior to their participation.

### Procedure

The study was carried out using a double-blind, sham tDCS-controlled, repeated measures, anodal stimulation paradigm. Testing was completed in brain stimulation dedicated laboratories within The School of Psychology at The University of Kent. Focal active (anodal) and placebo (sham) tDCS were administered in two sessions, with at least 72 hours between sessions. All participants were assessed for fluid intelligence using the matrix reasoning item bank (MaRs-IB; Chierchia et al., 2019) prior to the first stimulation session.

Participants also completed a short version of the object and action naming task for baseline performances (see Figure 1). The short version consisted of 10 object and 10 action images. The short version consisted of different images to those used during the stimulation sessions.

**Figure 1.**
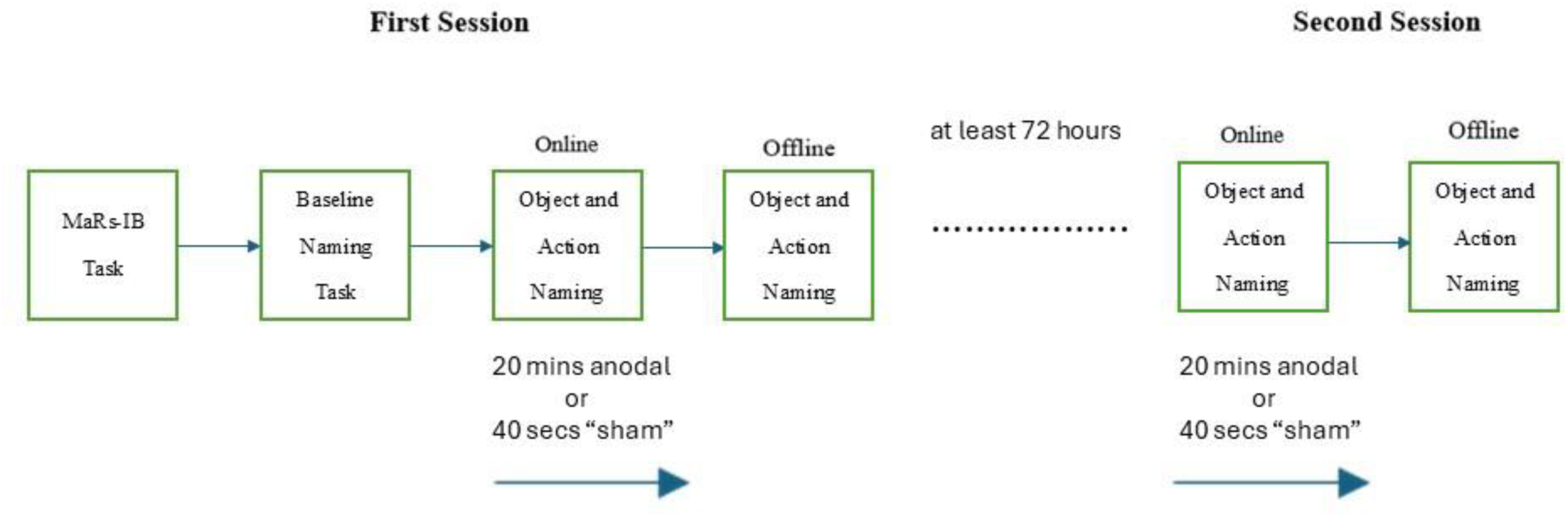
Study Design

In each session, we measured the participants’ naming performance twice, once whilst the participant was receiving stimulation (online) and once again after the stimulation had finished (offline), following the completion of the online stimulation. Four different matched sets of pictures were created and used for the online and offline sessions across the two testing sessions. The Black Box tool kit (https://www.blackboxtoolkit.com/urpvk.html) was used to record all responses and achieve millisecond precise response times.

### Fluid Intelligence

Fluid intelligence was assessed using MaRs-IB patterns (Chierchia et al., 2019). A single 3 x 3 matrix of patterns was presented with one missing tile. Participants were presented with four patterns and tasked with selecting the pattern that completed the matrix. The task is an open source alternative to abstract reasoning tasks that are often used to assess fluid intelligence, such as the matrix reasoning from standard IQ measures. Such tasks are strongly associated with Spearman’s *g* factor and show consistent age-related decline (Martin et al., 2021). Accuracy was summed across 40 trials and used as a measure of fluid intelligence. Five practice trials were completed prior to the test.

### Picture Naming Task

We acquired 200 black-and-white object drawings and 200 action drawings from the International Picture Naming Project (Bates et al., 2000) for use in the research. We created four sets of pictures (50 objects and 50 actions) equal in difficulty based on age of acquisition and frequency. A one-second fixation cross preceded each picture, followed by a bell sound which acted as a starting point for response time calculations. Each picture was presented for 3 seconds or until a verbal response was provided. Participants were instructed to name each picture as fast and accurate as possible. Object and action naming blocks were presented separately and breaks were offered between. Object naming blocks were always completed prior to action naming blocks. Semantically related words were marked as correct if deemed to accurately describe the image by a speech and language therapist (AY). Response times were our primary outcome and these were included only for correct responses. Accuracy was almost at ceiling throughout the study and was not considered for analysis. Figure 2 provides a schematic of the task procedure.

**Figure 2.**
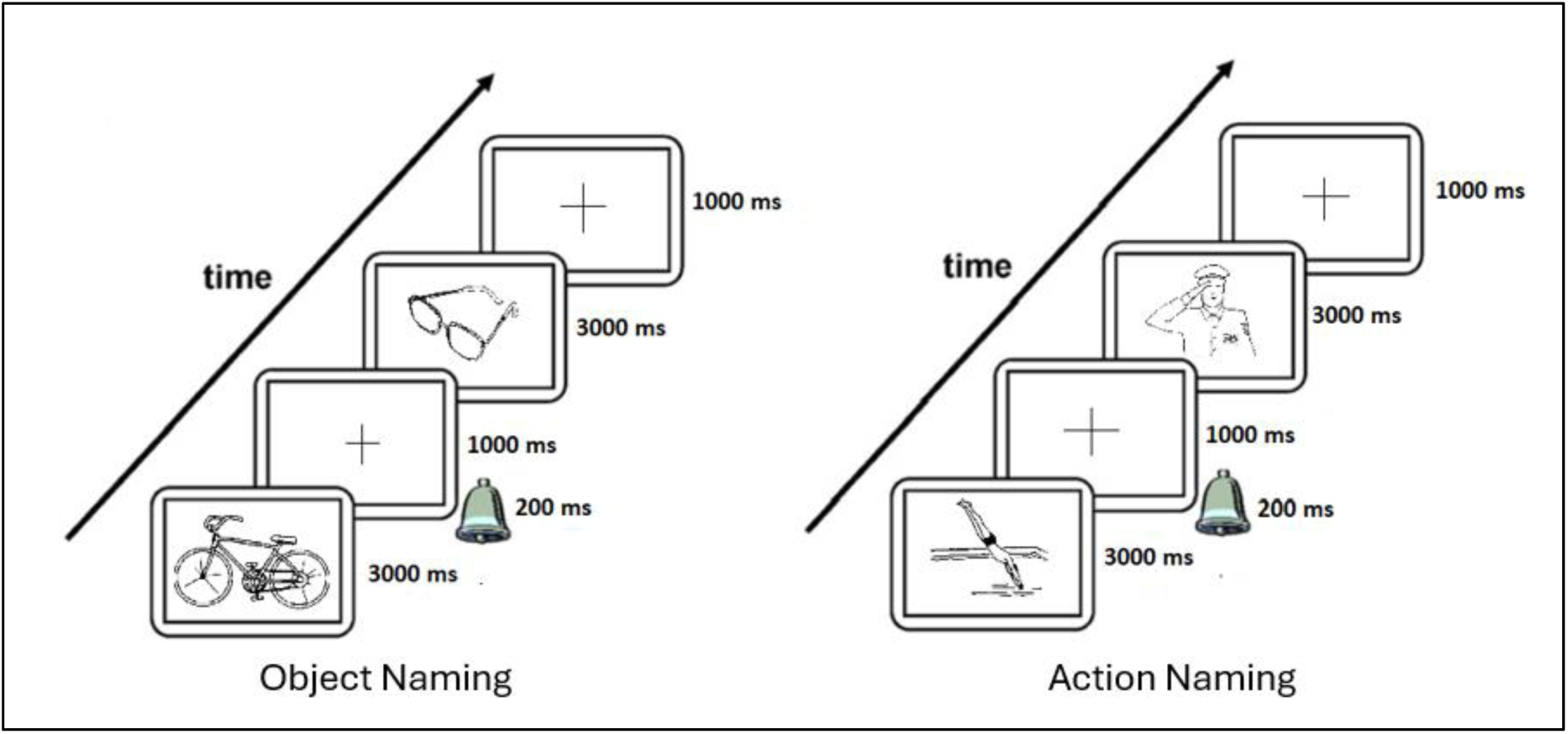
Picture Naming Task Procedure

### Focal transcranial direct current stimulation

We administered tDCS using a battery-driven, one-channel direct current stimulator (DC-Stimulator Plus, NeuroConn, Ilmenau, Germany). The focal tDCS electrode montage comprised two concentric conductive rubber electrodes, a small circular centre electrode (diameter = 2.5 cm), and a ring-shaped return electrode (inner diameter = 7.5 cm; outer diameter = 9.8 cm). The centre electrode was positioned over the left inferior frontal gyrus (position FC5 of the 10–20 EEG system) or left temporoparietal junction (position CP5 of the 10–20 EEG system). Current modelling for both the left IFG and right TPJ is provided in Figure 3. The modelling of current flow was based on a realistic 1 mm MNI152 T1 standard brain of a young adult. Current modelling was performed using ROAST (vOlumetric Approach to Simulate Transcranial Electric Stimulation), which is a fully automated, open-source MATLAB application (Huang et al., 2019). As no MRI data was available to run individualised modelling, the modelling presented in the current study should be considered as approximations of current flow based on a standard brain from a young adult.

**Figure 3.**
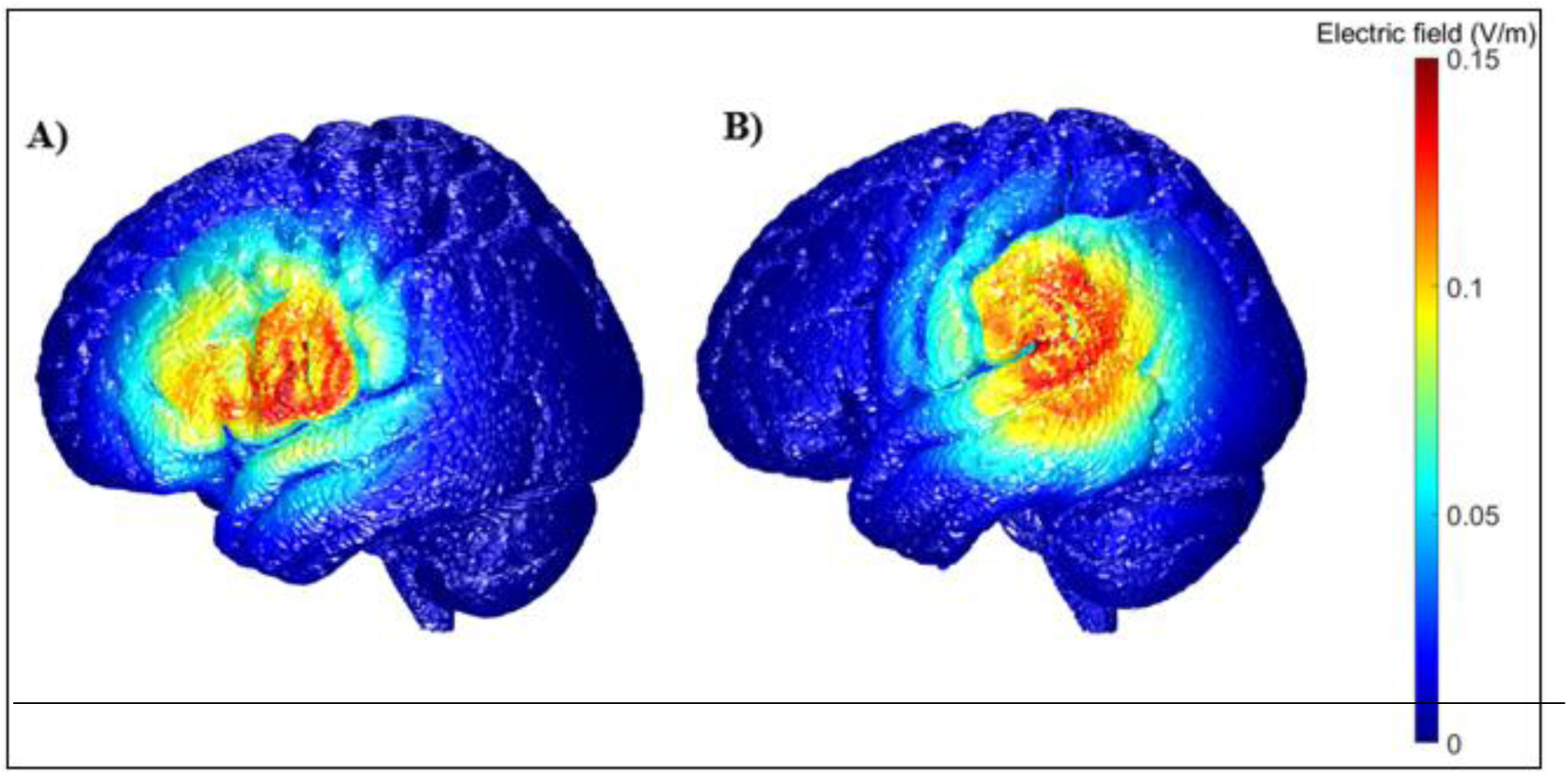
Surface rendering of the distribution and intensity of the estimated electric field for the two target sites with anodal stimulation. A) Left inferior frontal gyrus, B) Left temporoparietal junction.

The current was ramped up over 5 seconds to 1 mA during both stimulation conditions. In the anodal condition the current remained at 1mA for 20 mins before ramping down over 5 seconds. In the sham condition, the current ramped up to 1mA over 5 seconds and was ramped down after 40 seconds. The sham condition generates a tactile sensation comparable to the active stimulation. This approach has been shown to effectively blind subjects as to which stimulation they have received (Gbadeyan et al., 2016; Martin, Dzafic, et al., 2017; Martin et al., 2020; Martin, Huang, et al., 2019). We used the “study mode” of the DC stimulator, which involves a predefined code that triggers either active or sham tDCS, to achieve investigator blinding. The codes were assigned by a researcher who did not participate in data collection.

### Mood, adverse effects, and blinding

Participant’s mood was evaluated using the Visual Analogue Mood Scale (VAMS; Folstein & Luria, 1973) before and after each session. The VAMS assessed current positive and negative emotional states on visual analogue scales ranging from 0 to 100 (i.e., afraid, confused, sad, angry, energetic, tired, happy, and tense). Higher scores indicate greater intensity. Mood change scores were calculated by subtracting pre-stimulation scores from the post-stimulation scores for each mood. We calculated two mood change scores; positive and negative mood by summing the respective mood change scores.

Adverse effects were assessed using the self-report questionnaire developed by Brunoni et al. (2011). Participants graded the severity (1 = absent, 2 = mild, 3 = moderate, 4 = severe) and incidence of a variety of potential adverse effects, which consisted of headache, neck pain, scalp pain, tingling, itching, burning sensation, skin redness, sleepiness, trouble concentrating, and acute mood changes. Participant blinding was assessed at the completion of the study Participants were asked, “Do you think the active stimulation was in the first or second session?” Participants were forced to answer session one or two, even when they were unsure.

### Statistical Analyses

We conducted a 2×2×2×2× 2 repeated-measures ANOVA to assess whether online or offline stimulation to the lIFG or lTPJ modified object or action naming in healthy younger and older adults using a sham-controlled design. All factors were within-subject, except for age group and brain region, which were between-subject. The dependent variable was response speed.

In follow-up analyses, we explored whether baseline naming ability predicted stimulation response in older adults. A 2×2×2×2 repeated-measures ANCOVA was conducted with baseline naming and fluid intelligence as covariates. The same analysis was also conducted in younger adults.

We assessed blinding across each age group and stimulation site using chi-square tests. Mood change was assessed by calculating the difference on a combined positive scale (happy and energetic) and combined negative scales (tired, afraid, sad, tense, confused, angry) of the VAMS before and after both anodal and sham stimulation. The positive and negative mood change scores were the dependent variables in 2×2×2 ANOVAs, with age group (young vs. older), stimulation site (lIFG vs. lTPJ), and stimulation type (sham vs. anodal) as independent variables.

## Results

A main effect of **Naming Type** was identified, *F*(1, 140) = 1607.87, *p* < .001, η²ₚ = 0.92, but this was subsumed within a significant **Age Group × Naming Type** interaction, *F*(1, 140) = 24.31, *p* < .001, η²ₚ = 0.15. The post hoc analysis identified slower action naming in younger adults (*M* = 1380 ms) than in older adults (*M* = 1294 ms). No difference was identified for object naming in younger adults (*M* = 1022 ms) and older adults (*M* = 1014 ms).

The main effect of **Stimulation** was not significant, *F*(1, 140) = 0.37, *p* = .54, η²ₚ = 0.003. Likewise, stimulation had no effect on performance dependent upon **Region, Age Group, Stimulation Time**, or **Naming Type**, nor any interaction between these variables (*p*s between .09 and .99; see Appendix 1 for full details).

## Stimulation effects dependent on baseline functioning

### Baseline Naming Speed

We demonstrated that the interaction between **Baseline Naming × Stimulation Type × Region** was significant, *F*(1, 66) = 4.41, *p* = .04, η²ₚ = 0.06. However, the **Fluid Intelligence × Stimulation Type × Region** interaction was not significant, *F*(1, 66) = 0.38, *p* = .54, η²ₚ = 0.01. Further interactions, including **Baseline Naming × Stimulation Type × Region × Stimulation Time**, *F*(1, 68) = 0.34, *p* = .56, η²ₚ = 0.001, and **Baseline Naming × Stimulation Type × Region × Stimulation Time × Naming Type**, *F*(1, 68) = 0.08, *p* = .77, η²ₚ = 0.001, were not significant. The full statistical model is provided in Appendix 2.

To explore the significant interaction, we conducted separate analyses for each brain region. For the lIFG, the **Baseline Naming × Stimulation Type** interaction was significant, *F*(1, 33) = 6.74, *p* = .01, η²ₚ = 0.17. For the lTPJ, this interaction was not significant, *F*(1, 33) = 0.65, *p* = .43, η²ₚ = 0.02. Scatterplots (see Figure 4) showed that stimulation response was negatively correlated with baseline naming performance. Therefore, stimulation to the lIFG in older adults increased naming speed in those who performed slower during the baseline assessment.

**Figure 4.**
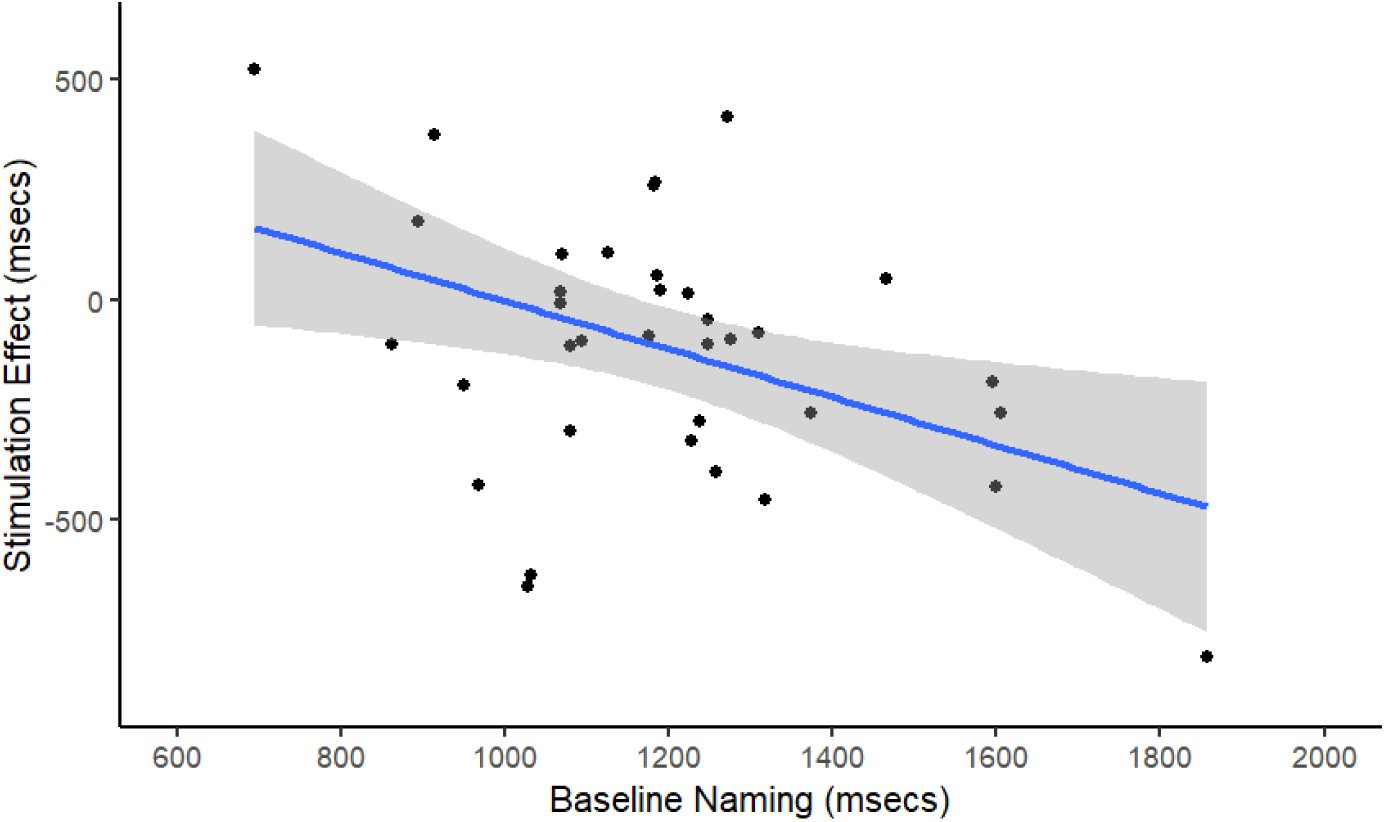
Stimulation effect at the left inferior frontal gyrus was dependent on baseline naming speed

We conducted the same analysis in younger adults, but no relationship was identified for baseline naming or fluid intelligence on any of the stimulation effects (*p*s between .11 and .91; see Appendix 3 for more details).

### Blinding

We used a chi-squared test to investigate how participants’ age group and stimulated region affected their ability to properly identify the active session, demonstrating the efficiency of blinding. The chi-squared test revealed a significant association between age group and blinding scores, χ²(1, N = 64) = 6.399, *p* = .011, suggesting that age group influenced participants’ ability to maintain blinding integrity. Younger adults (24/32 correct at lIFG; 25/32 correct at lTPJ) were able to guess active sessions better than older adults (16/32 correct at lIFG; 18/32 correct at lTPJ). Blinding efficacy was comparable across the two stimulation sites, χ²(1, N = 64) = 0.256, *p* = .61.

### Positive Mood

We conducted a 2×2×2 repeated-measures ANOVA to determine whether Anodal and Sham sessions with older and younger adults at the left IFG and TPJ influenced VAMS positive mood scores. A significant stimulation effect was found, *F*(1, 140) = 4.508, *p* = .035, η²ₚ = 0.012. Participants reported a small but significant reduction in VAMS positive scores during the Anodal stimulation session (*M* = −9.996) compared to the Sham session (*M* = −2.461). No interactions were identified for Stimulation × Age Group, *F*(1, 140) = 0.014, *p* = .907, η²ₚ < 0.001; Stimulation × Region, *F*(1, 140) = 0.074, *p* = .785; or Stimulation × Age Group × Region, *F*(1, 140) = 3.511, *p* = .063, η²ₚ = 0.02. Change scores are presented in Table 2.

**Table 1.**
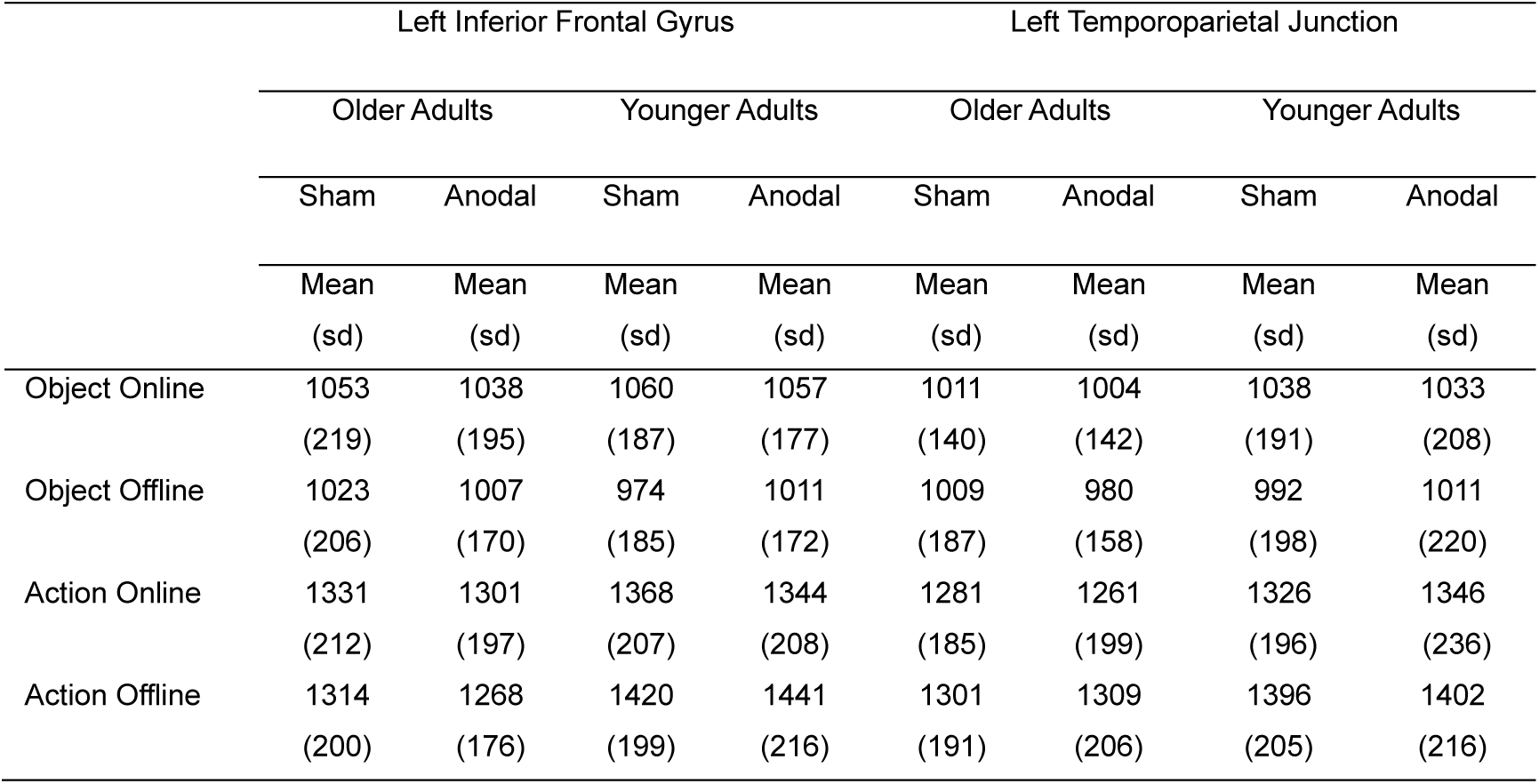
Descriptive statistics of performance (response times in milliseconds) on object and action naming during the online and offline phases of the stimulation across the stimulation site, age group, and stimulation type.

|  | Left Inferior Frontal Gyrus |  |  |  | Left Temporoparietal Junction |  |  |  |
| --- | --- | --- | --- | --- | --- | --- | --- | --- |
|  | Older Adults |  | Younger Adults |  | Older Adults |  | Younger Adults |  |
|  | Sham | Anodal | Sham | Anodal | Sham | Anodal | Sham | Anodal |
|  | Mean<br>(sd) | Mean<br>(sd) | Mean<br>(sd) | Mean<br>(sd) | Mean<br>(sd) | Mean<br>(sd) | Mean<br>(sd) | Mean<br>(sd) |
| Object Online | 1053<br>(219) | 1038<br>(195) | 1060<br>(187) | 1057<br>(177) | 1011<br>(140) | 1004<br>(142) | 1038<br>(191) | 1033<br>(208) |
| Object Offline | 1023<br>(206) | 1007<br>(170) | 974<br>(185) | 1011<br>(172) | 1009<br>(187) | 980<br>(158) | 992<br>(198) | 1011<br>(220) |
| Action Online | 1331<br>(212) | 1301<br>(197) | 1368<br>(207) | 1344<br>(208) | 1281<br>(185) | 1261<br>(199) | 1326<br>(196) | 1346<br>(236) |
| Action Offline | 1314<br>(200) | 1268<br>(176) | 1420<br>(199) | 1441<br>(216) | 1301<br>(191) | 1309<br>(206) | 1396<br>(205) | 1402<br>(216) |

**Table 2.** Change in mood scores for sham and anodal stimulation across regions and age groups.

|  | IIFG |  |  |  | ITPJ |  |  |  |
| --- | --- | --- | --- | --- | --- | --- | --- | --- |
|  | Older Adults |  | Younger Adults |  | Older Adults |  | Younger Adults |  |
|  | Sham | Anodal | Sham | Anodal | Sham | Anodal | Sham | Anodal |
|  | Mean<br>(sd) | Mean<br>(sd) | Mean<br>(sd) | Mean<br>(sd) | Mean<br>(sd) | Mean<br>(sd) | Mean<br>(sd) | Mean<br>(sd) |
| Change in Positive Mood | -0.41<br>(41.22) | -0.75<br>(40.37) | 0.33<br>(32.81) | -12.75<br>(38.16) | -1.91<br>(39.57) | -17.82<br>(41.39) | -7.19<br>(15.98) | -8.66<br>(21.92) |
| Change in Negative Mood | -20.58<br>(41.29) | -12.65<br>(78.17) | -9.32<br>(50.36) | -16.35<br>(45.48) | -20.59<br>(42.05) | -25.27<br>(46.15) | -11<br>(30.47) | -11.72<br>(51.89) |

IIFG = left inferior frontal gyrus; ITPJ = left temporoparietal junction; sd = standard deviation.

### Negative Mood

No effect of stimulation on VAMS negative scores was observed, *F*(1, 140) = 1.033, *p* = .311, η²ₚ = 0.002. Stimulation also did not affect VAMS negative scores for the following interactions: Stimulation × Age Group, *F*(1, 140) = 0.031, *p* = .860, η²ₚ < 0.001; Stimulation × Region, *F*(1, 140) = 0.186, *p* = .667, η²ₚ = 0.001; or Stimulation × Age Group × Region, *F*(1, 140) = 0.063, *p* = .802, η²ₚ < 0.001. Change scores are presented in Table 2.

### Adverse Effects

Participants reported slightly higher mild adverse effect scores for the Anodal session (*M* = 1.826) compared to the Sham session (*M* = 1.556; see Table 3 for details). However, the stimulation effect was not significant, *F*(1, 140) = 3.283, *p* = .072, η²ₚ = 0.02. No interaction effects were identified for Stimulation × Age Group, *F*(1, 140) = 0.486, *p* = .487, η²ₚ = 0.003; Stimulation × Region, *F*(1, 140) = 0.002, *p* = .963, η²ₚ < 0.001; or Stimulation × Age Group × Region, *F*(1, 140) = 0.779, *p* = .379, η²ₚ = 0.006.

**Table 3.** Adverse effects across stimulation site, age group, and stimulation type.

|  | IIFG |  |  |  | ITPJ |  |  |  |
| --- | --- | --- | --- | --- | --- | --- | --- | --- |
|  | Older Adults |  | Younger Adults |  | Older Adults |  | Younger Adults |  |
|  | Sham | Anodal | Sham | Anodal | Sham | Anodal | Sham | Anodal |
|  | Mean<br>(sd) | Mean<br>(sd) | Mean<br>(sd) | Mean<br>(sd) | Mean<br>(sd) | Mean<br>(sd) | Mean<br>(sd) | Mean<br>(sd) |
| Headache | 1<br>(0) | 1.06<br>(0.33) | 1.11<br>(0.32) | 1.17<br>(0.45) | 1.08<br>(0.28) | 1.06<br>(0.23) | 1.31<br>(0.52) | 1.20<br>(0.47) |
| Neck Pain | 1.05<br>(0.23) | 1.02<br>(0.17) | 1.06<br>(0.33) | 1.06<br>(0.23) | 1.06<br>(0.33) | 1.08<br>(0.37) | 1.03<br>(0.17) | 1.03<br>(0.17) |
| Scalp Pain | 1.03<br>(0.17) | 1.06<br>(0.23) | 1.14<br>(0.42) | 1.08<br>(0.29) | 1.06<br>(0.33) | 1.14<br>(0.42) | 1.03<br>(0.17) | 1.14<br>(0.42) |
| Tingling | 1.36<br>(0.49) | 1.08<br>(0.50) | 1.42<br>(0.65) | 1.61<br>(0.64) | 1.53<br>(0.70) | 1.55<br>(0.70) | 1.77<br>(0.77) | 1.97<br>(0.88) |
| Itching | 1.06<br>(0.23) | 1.03<br>(0.17) | 1.17<br>(0.51) | 1.19<br>(0.40) | 1.08<br>(0.37) | 1.14<br>(0.49) | 1.14<br>(0.35) | 1.25<br>(0.50) |
| Burning | 1<br>(0) | 1.06<br>(0.24) | 1.08<br>(0.37) | 1.25<br>(0.55) | 1.17<br>(0.61) | 1.11<br>(0.40) | 1.17<br>(0.38) | 1.28<br>(0.70) |
| Skin Redness | 1<br>(0) | 1<br>(0) | 1.05<br>(0.33) | 1<br>(0) | 1<br>(0) | 1<br>(0) | 1.03<br>(0.17) | 1.08<br>(0.28) |
| Sleepiness | 1.08<br>(0.28) | 1.06<br>(0.23) | 1.44<br>(0.61) | 1.50<br>(0.61) | 1.19<br>(0.40) | 1.14<br>(0.35) | 1.53<br>(0.74) | 1.47<br>(0.61) |
| Trouble Concentrating | 1.06<br>(0.23) | 1.14<br>(0.42) | 1.31<br>(0.52) | 1.19<br>(0.40) | 1.28<br>(0.51) | 1.25<br>(0.44) | 1.28<br>(0.45) | 1.33<br>(0.63) |
| Mood change | 1.03<br>(0.17) | 1.06<br>(0.23) | 1.06<br>(0.23) | 1.03<br>(0.17) | 1<br>(0) | 1<br>(0) | 1<br>(0) | 1.03<br>(0.17) |

IIFG = left inferior frontal gyrus; ITPJ = left temporoparietal junction; sd = standard deviation

## Discussion

We explored the effect of online and offline focal stimulation to either the left inferior frontal gyrus (lIFG) or left temporoparietal junction (lTPJ) on object and action naming in a large sample of young and older adults. Moreover, we assessed whether baseline cognitive performance was a predictor of stimulation response in the older adults. We demonstrated that focal stimulation to the lIFG, but not the lTPJ, can improve both object and action naming in older adults, but crucially, only in those who show lower baseline naming performance. Therefore, our results demonstrate that tDCS can improve language performance in those that require a greater “boost”. The study builds on existing evidence that tDCS is most beneficial for those with lower cognitive function (Learmonth et al., 2015; Meinzer et al., 2016; Perceval et al., 2020). In the absence of an overall effect of tDCS (across the entire group), this suggests that inconsistent effects reported in the literature (Koo et al., 2023), may be due to the failure to consider intraindividual differences in baseline performance. We also demonstrate no difference in online or offline stimulation effects, in young or older adults, but do show site-specific effects at the lIFG. Older adults only identified active vs. sham tDCS at chance level, making it unlikely that stimulation effects in this group are explained by unblinding. It was of interest that the younger adults were better able to guess the stimulation condition at both lIFG and lTPJ for the focal stimulation protocol. Similar findings have been demonstrated using conventional tDCS (Gandiga et al., 2006; Kessler et al., 2012) and likely reflect age-related changes to skin conductance and the density or effectiveness of small fiber nerve endings. In line with the vast majority of tDCS studies that used conventional tDCS (Antal et al. 2017), our focal set-up only induced mild adverse effects. This confirms the safety of this approach, thereby rendering it as an attractive approach as neuroenhancement technique in advanced age and age-related neurodegenerative conditions (Kortteenniemi et al., 2017).

Baseline dependent effects align with previous studies focused on associative learning (Perceval et al., 2020), visual detection (Learmonth et al., 2015), processing speed (Riemann, Mittelstädt, et al., 2024), and word generation (Meinzer et al., 2013), but the current study is the first to show such effects on picture naming. Moreover, we show that baseline performance should be measured in relation to the task studied, as general cognitive ability, as measured using matrix reasoning, had no influence on stimulation response. Therefore, stimulation may act on local neural inefficiencies related to naming, rather than other broad measures of brain function, such as fluid intelligence, which shows consistent age-related decline (Bugg et al., 2006), and likely reflects neural changes in a broad multiple demand network of fronto-parietal activity, separate from language systems (Woolgar et al., 2018). The results highlight the need to assess specific cognitive function to determine likely benefits from tDCS, with clear implications for personalising brain stimulation protocols in healthy ageing and in pathological ageing conditions.

Our results show baseline performance dependent effects of focal stimulation to the lIFG only, with stimulation to the lTPJ having no effect. Faster picture naming due to lIFG stimulation was identified for both object and action words, providing evidence against our tentative hypothesis of stronger frontal effects for action naming. This is inconsistent with previous tDCS studies showing specific effects on action naming following stimulation to the lIFG and object naming following stimulation to the lTPJ. However, previous effects were observed in studies with much smaller samples and used fewer experimental stimuli (Fiori et al., 2011, 2013; Marangolo et al., 2013). Moreover, our results are consistent with previous theoretical accounts positing that a dissociation in relation to grammatical class is not observed at the lIFG (Scott, 2006). Stimulation to the lTPJ has generally resulted in weaker effects on language functions, even with focal stimulation (Perceval et al., 2017; Riemann, van Lück, et al., 2024). Moreover, previous studies using conventional tDCS may actually be stimulating neighbouring neural regions such as the motor cortex, middle, or superior temporal gyrus, with little stimulation to the lTPJ (Riemann, van Lück, et al., 2024). Our results suggest that targeting the lIFG is optimal for improving language function in older adults.

Although the results of the current study are encouraging, further research is required to determine the efficacy of tDCS in clinical groups associated with impaired naming performance or anomia, such as aphasia, semantic dementia, or Parkinson’s Disease (Laine & Martin, 2006; Verhaegen et al., 2022; Woollams et al., 2008). It is also important to investigate how long such improvements to naming speed last and whether multisession tDCS can result in long-lasting improvements to language, increasing its attractiveness for clinical use. Previous studies using a multisession approach have shown improvements lasting 3 months in older adults with lower baseline cognitive levels (Perceval et al., 2020) in a protocol assessing associative learning. The results provide encouragement for further research investigating multisession effects on naming speed, and language more broadly, in older adults. The effects of baseline cognitive performance would likely be enhanced with a more representative sample of older adults across the cognitive spectrum. Effects may also be enhanced through a more personalised approach to locating and targeting the lIFG via structural imaging and neuronavigation and affixing electrodes accordingly. In the current study we used a standard approach based on EEG coordinates, but this fails to consider interindividual variability. Moreover, subtle electrode placement errors have been shown to affect current flow to a greater extent when using focal stimulation methods (Niemann, Riemann, et al., 2024) and future research employing a more personalised neuronavigated approach (Niemann, Shababaie, et al., 2024) or a comparison with standard tDCS approaches, is required. Importantly, because this study had no access to MRI images for individualized modelling, the current modelling provided reflects an estimate of electric fields based on a standardised, young structural template. Future research should include individualised structural mapping to understand age-related differences in current flow and individual differences in relation to stimulation effects.

The current study provides novel evidence regarding the timing of stimulation effects, with both online and offline stimulation improving naming speed in lower functioning older adults. The results suggest that inconsistencies reported previously cannot solely be explained by the timing of stimulation protocols. Future research could provide more comprehensive evidence by conducting a between-subjects analysis of stimulation timing to avoid potential interactions between stimulation and familiarity of task or training which is unable to separated using our design.

In sum, we have demonstrated that focal tDCS to the lIFG can improve naming speed in older adults with lower baseline naming ability. Fluid intelligence did not predict tDCS effects, which highlights the task specificity of this effect . We found no differences between online and offline stimulation suggesting either protocol improves performance in lower functioning older adults. Stimulation effects were limited to the lIFG, but comparable for object and action naming. Improving performance in lower functioning adults increases the evidence that tDCS may be a suitable clinical tool for populations with impairments to language in age-related neurodegenerative conditions.

## Acknowledgements

We would like to acknowledge all participants for their time and effort.

