## Appendix 1 for "Regionally specific picture naming benefits of focal tDCS are dependent on baseline performance in older adults"

**Appendix 1. Full model statistical output**

| **Within Subjects Effects** | | | | | | | | | | | | | | |
| --- | --- | --- | --- | --- | --- | --- | --- | --- | --- | --- | --- | --- | --- | --- |
| **Cases** | **Sum of Squares** | | | **df** | | **Mean Square** | | | **F** | | **p** | | **η²_p_** | |
| Stimulation Type |  | 0.008 |  | | 1 |  | 0.008 |  | | 0.369 |  | 0.545 |  | 0.003 |
| Stimulation Type ✻ Region |  | 0.005 |  | | 1 |  | 0.005 |  | | 0.242 |  | 0.624 |  | 0.002 |
| Stimulation Type ✻ Age Group |  | 0.056 |  | | 1 |  | 0.056 |  | | 2.766 |  | 0.099 |  | 0.019 |
| Stimulation Type ✻ Region ✻ Age Group |  | 0.002 |  | | 1 |  | 0.002 |  | | 0.121 |  | 0.728 |  | <0.001 |
| Residuals |  | 2.846 |  | | 140 |  | 0.020 |  | |  |  |  |  |  |
| Naming Type |  | 29.351 |  | | 1 |  | 29.351 |  | | 1607.871 |  | < .001 |  | 0.920 |
| Naming Type ✻ Region |  | <0.001 |  | | 1 |  | <0.001 |  | | 0.024 |  | 0.877 |  | <0.001 |
| Naming Type ✻ Age Group |  | 0.444 |  | | 1 |  | 0.444 |  | | 24.306 |  | < .001 |  | 0.148 |
| Naming Type ✻ Region ✻ Age Group |  | 0.019 |  | | 1 |  | 0.019 |  | | 1.066 |  | 0.304 |  | 0.008 |
| Residuals |  | 2.556 |  | | 140 |  | 0.018 |  | |  |  |  |  |  |
| Stimulation Time |  | <0.001 |  | | 1 |  | <0.001 |  | | 0.003 |  | 0.954 |  | <0.001 |
| Stimulation Time ✻ Region |  | 0.042 |  | | 1 |  | 0.042 |  | | 4.420 |  | 0.037 |  | 0.031 |
| Stimulation Time ✻ Age Group |  | 0.023 |  | | 1 |  | 0.023 |  | | 2.398 |  | 0.124 |  | 0.017 |
| Stimulation Time ✻ Region ✻ Age Group |  | 0.014 |  | | 1 |  | 0.014 |  | | 1.466 |  | 0.228 |  | 0.010 |
| Residuals |  | 1.341 |  | | 140 |  | 0.010 |  | |  |  |  |  |  |
| Stimulation Type ✻ Naming Type |  | 0.002 |  | | 1 |  | 0.002 |  | | 0.238 |  | 0.626 |  | 0.002 |
| Stimulation Type ✻ Naming Type ✻ Region |  | 0.015 |  | | 1 |  | 0.015 |  | | 1.677 |  | 0.197 |  | 0.012 |
| Stimulation Type ✻ Naming Type ✻ Age Group |  | <0.001 |  | | 1 |  | <0.001 |  | | 0.005 |  | 0.946 |  | <0.001 |
| Stimulation Type ✻ Naming Type ✻ Region ✻ Age Group |  | <0.001 |  | | 1 |  | <0.001 |  | | 0.063 |  | 0.802 |  | <0.001 |
| Residuals |  | 1.277 |  | | 140 |  | 0.009 |  | |  |  |  |  |  |
| Stimulation Type ✻ Stimulation Time |  | 0.008 |  | | 1 |  | 0.008 |  | | 0.958 |  | 0.329 |  | 0.007 |
| Stimulation Type ✻ Stimulation Time ✻ Region |  | 0.003 |  | | 1 |  | 0.003 |  | | 0.362 |  | 0.548 |  | 0.003 |
| Stimulation Type ✻ Stimulation Time ✻ Age Group |  | 0.013 |  | | 1 |  | 0.013 |  | | 1.666 |  | 0.199 |  | 0.012 |
| Stimulation Type ✻ Stimulation Time ✻ Region ✻ Age Group |  | 0.011 |  | | 1 |  | 0.011 |  | | 1.406 |  | 0.238 |  | 0.010 |
| Residuals |  | 1.113 |  | | 140 |  | 0.008 |  | |  |  |  |  |  |
| Naming Type ✻ Stimulation Time |  | 0.378 |  | | 1 |  | 0.378 |  | | 70.427 |  | < .001 |  | 0.335 |
| Naming Type ✻ Stimulation Time ✻ Region |  | <0.001 |  | | 1 |  | <0.001 |  | | 0.011 |  | 0.918 |  | <0.001 |
| Naming Type ✻ Stimulation Time ✻ Age Group |  | 0.153 |  | | 1 |  | 0.153 |  | | 28.456 |  | < .001 |  | 0.169 |
| Naming Type ✻ Stimulation Time ✻ Region ✻ Age Group |  | 0.033 |  | | 1 |  | 0.033 |  | | 6.150 |  | 0.014 |  | 0.042 |
| Residuals |  | 0.751 |  | | 140 |  | 0.005 |  | |  |  |  |  |  |
| Stimulation Type ✻ Naming Type ✻ Stimulation Time |  | <0.001 |  | | 1 |  | <0.001 |  | | <0.001 |  | 0.987 |  | <0.001 |
| Stimulation Type ✻ Naming Type ✻ Stimulation Time ✻ Region |  | <0.001 |  | | 1 |  | <0.001 |  | | 0.065 |  | 0.799 |  | <0.001 |
| Stimulation Type ✻ Naming Type ✻ Stimulation Time ✻ Age Group |  | 0.005 |  | | 1 |  | 0.005 |  | | 0.561 |  | 0.455 |  | 0.004 |
| Stimulation Type ✻ Naming Type ✻ Stimulation Time ✻ Region ✻ Age Group |  | 0.013 |  | | 1 |  | 0.013 |  | | 1.494 |  | 0.224 |  | 0.011 |
| Residuals |  | 1.244 |  | | 140 |  | 0.009 |  | |  |  |  |  |  |
| *Note.*  Type III Sum of Squares | | | | | | | | | | | | | | |

| **Between Subjects Effects** | | | | | | | | | | | | |
| --- | --- | --- | --- | --- | --- | --- | --- | --- | --- | --- | --- | --- |
| **Cases** | | **Sum of Squares** | | **df** | | **Mean Square** | | **F** | | **p** | | **η²_p_** |
| Region |  | 0.110 |  | 1 |  | 0.110 |  | 0.490 |  | 0.485 |  | 0.003 |
| Age Group |  | 0.590 |  | 1 |  | 0.590 |  | 2.616 |  | 0.108 |  | 0.018 |
| Region ✻ Age Group |  | 0.003 |  | 1 |  | 0.003 |  | 0.014 |  | 0.908 |  | 9.667×10^-5^ |
| Residuals |  | 31.570 |  | 140 |  | 0.226 |  |  |  |  |  |  |
| *Note.*  Type III Sum of Squares | | | | | | | | | | | | |
