## Appendix 2 for "Regionally specific picture naming benefits of focal tDCS are dependent on baseline performance in older adults"

**Appendix 2. Effects of baseline naming and fluid intelligence on stimulation effects in older adults**

| **Cases** | **Sum of Squares** | **df** | **Mean Square** | **F** | **p** | **η²p** |
| --- | --- | --- | --- | --- | --- | --- |
| Stimulation Type | < .001 | 1 | < .001 | 0.001 | 0.977 | < .001 |
| Stimulation Type ✻ Region | 0.014 | 1 | 0.014 | 1.111 | 0.296 | 0.017 |
| Stimulation Type ✻ Fluid Intelligence | 0.003 | 1 | 0.003 | 0.206 | 0.651 | 0.003 |
| Stimulation Type ✻ Baseline Naming Speed | 0.006 | 1 | 0.006 | 0.472 | 0.494 | 0.007 |
| Stimulation Type ✻ Region ✻ Baseline Naming Speed | 0.054 | 1 | 0.054 | 4.406 | 0.040 | 0.063 |
| Stimulation Type ✻ Region ✻ Fluid Intelligence | 0.005 | 1 | 0.005 | 0.378 | 0.541 | 0.006 |
| Residuals | 0.807 | 66 | 0.012 |  |  |  |
| Naming Type | 0.022 | 1 | 0.022 | 1.178 | 0.282 | 0.018 |
| Naming Type ✻ Region | 0.046 | 1 | 0.046 | 2.387 | 0.127 | 0.035 |
| Naming Type ✻ Fluid Intelligence | 0.003 | 1 | 0.003 | 0.172 | 0.680 | 0.003 |
| Naming Type ✻ Baseline Naming Speed | 0.087 | 1 | 0.087 | 4.557 | 0.037 | 0.065 |
| Naming Type ✻ Region ✻ Baseline Naming Speed | 0.089 | 1 | 0.089 | 4.674 | 0.034 | 0.066 |
| Naming Type ✻ Region ✻ Fluid Intelligence | < .001 | 1 | < .001 | 0.007 | 0.936 | < .001 |
| Residuals | 1.258 | 66 | 0.019 |  |  |  |
| Stimulation Time | 0.002 | 1 | 0.002 | 0.169 | 0.682 | 0.003 |
| Stimulation Time ✻ Region | 0.011 | 1 | 0.011 | 1.130 | 0.292 | 0.017 |
| Stimulation Time ✻ Fluid Intelligence | < .001 | 1 | < .001 | 0.045 | 0.834 | < .001 |
| Stimulation Time ✻ Baseline Naming Speed | 0.008 | 1 | 0.008 | 0.819 | 0.369 | 0.012 |
| Stimulation Time ✻ Region ✻ Baseline Naming Speed | < .001 | 1 | < .001 | 0.033 | 0.856 | < .001 |
| Stimulation Time ✻ Region ✻ Fluid Intelligence | 0.024 | 1 | 0.024 | 2.469 | 0.121 | 0.036 |
| Residuals | 0.640 | 66 | 0.010 |  |  |  |
| Stimulation Type ✻ Naming Type | 0.001 | 1 | 0.001 | 0.141 | 0.709 | 0.002 |
| Stimulation Type ✻ Naming Type ✻ Region | 0.010 | 1 | 0.010 | 1.229 | 0.272 | 0.018 |
| Stimulation Type ✻ Naming Type ✻ Fluid Intelligence | 0.007 | 1 | 0.007 | 0.875 | 0.353 | 0.013 |
| Stimulation Type ✻ Naming Type ✻ Baseline Naming Speed | < .001 | 1 | < .001 | 0.038 | 0.846 | < .001 |
| Stimulation Type ✻ Naming Type ✻ Region ✻ Baseline Naming Speed | 0.003 | 1 | 0.003 | 0.342 | 0.561 | 0.005 |
| Stimulation Type ✻ Naming Type ✻ Region ✻ Fluid Intelligence | 0.007 | 1 | 0.007 | 0.886 | 0.350 | 0.013 |
| Residuals | 0.539 | 66 | 0.008 |  |  |  |
| Stimulation Type ✻ Stimulation Time | 0.003 | 1 | 0.003 | 0.411 | 0.523 | 0.006 |
| Stimulation Type ✻ Stimulation Time ✻ Region | 0.002 | 1 | 0.002 | 0.254 | 0.616 | 0.004 |
| Stimulation Type ✻ Stimulation Time ✻ Fluid Intelligence | < .001 | 1 | < .001 | 0.092 | 0.763 | < .001 |
| Stimulation Type ✻ Stimulation Time ✻ Baseline Naming Speed | 0.003 | 1 | 0.003 | 0.365 | 0.548 | 0.006 |
| Stimulation Type ✻ Stimulation Time ✻ Region ✻ Baseline Naming Speed | 0.003 | 1 | 0.003 | 0.383 | 0.538 | 0.006 |
| Stimulation Type ✻ Stimulation Time ✻ Region ✻ Fluid Intelligence | < .001 | 1 | < .001 | 0.031 | 0.861 | < .001 |
| Residuals | 0.540 | 66 | 0.008 |  |  |  |
| Naming Type ✻ Stimulation Time | 0.003 | 1 | 0.003 | 0.650 | 0.423 | 0.010 |
| Naming Type ✻ Stimulation Time ✻ Region | < .001 | 1 | < .001 | 0.056 | 0.814 | < .001 |
| Naming Type ✻ Stimulation Time ✻ Fluid Intelligence | 0.010 | 1 | 0.010 | 2.035 | 0.158 | 0.030 |
| Naming Type ✻ Stimulation Time ✻ Baseline Naming Speed | < .001 | 1 | < .001 | 0.157 | 0.693 | < .001 |
| Naming Type ✻ Stimulation Time ✻ Region ✻ Baseline Naming Speed | < .001 | 1 | < .001 | 0.003 | 0.953 | < .001 |
| Naming Type ✻ Stimulation Time ✻ Region ✻ Fluid Intelligence | < .001 | 1 | < .001 | 0.015 | 0.903 | < .001 |
| Residuals | 0.340 | 66 | 0.005 |  |  |  |
| Stimulation Type ✻ Naming Type ✻ Stimulation Time | 0.007 | 1 | 0.007 | 0.888 | 0.349 | 0.013 |
| Stimulation Type ✻ Naming Type ✻ Stimulation Time ✻ Region | < .001 | 1 | < .001 | 0.080 | 0.778 | < .001 |
| Stimulation Type ✻ Naming Type ✻ Stimulation Time ✻ Fluid Intelligence | 0.008 | 1 | 0.008 | 0.974 | 0.327 | 0.015 |
| Stimulation Type ✻ Naming Type ✻ Stimulation Time ✻ Baseline Naming Speed | 0.001 | 1 | 0.001 | 0.180 | 0.673 | 0.003 |
| Stimulation Type ✻ Naming Type ✻ Stimulation Time ✻ Region ✻ Baseline Naming Speed | < .001 | 1 | < .001 | 0.047 | 0.829 | < .001 |
| Stimulation Type ✻ Naming Type ✻ Stimulation Time ✻ Region ✻ Fluid Intelligence | 0.008 | 1 | 0.008 | 0.975 | 0.327 | 0.015 |
| Residuals | 0.542 | 66 | 0.008 |  |  |  |

*Note.*  Type III Sum of Square

**Between Subjects Effects**

| **Cases** | **Sum of Squares** | **df** | **Mean Square** | **F** | **p** | **η²p** |
| --- | --- | --- | --- | --- | --- | --- |
| Region | 0.001 | 1 | 0.001 | 0.008 | 0.930 | < .001 |
| Fluid Intelligence | 0.192 | 1 | 0.192 | 1.992 | 0.163 | 0.029 |
| Baseline Naming Speed | 7.253 | 1 | 7.253 | 75.401 | < .001 | 0.533 |
| Region ✻ Baseline Naming Speed | < .001 | 1 | < .001 | 0.004 | 0.948 | < .001 |
| Region ✻ Fluid Intelligence | < .001 | 1 | < .001 | < .001 | 0.987 | < .001 |
| Residuals | 6.349 | 66 | 0.096 |  |  |  |

*Note.*  Type III Sum of Square

**Descriptives**

| **Stimulation Type** | **Naming Type** | **Stimulation Time** | **Region** | **N** | **Mean** | **SD** | **SE** | **Coefficient of Variation** |
| --- | --- | --- | --- | --- | --- | --- | --- | --- |
| Sham | Object | Online | F | 36 | 1.053 | 0.219 | 0.037 | 0.208 |
|  |  |  | T | 36 | 1.011 | 0.140 | 0.023 | 0.138 |
|  |  | Offline | F | 36 | 1.023 | 0.206 | 0.034 | 0.201 |
|  |  |  | T | 36 | 1.009 | 0.187 | 0.031 | 0.185 |
|  | Action | Online | F | 36 | 1.331 | 0.212 | 0.035 | 0.160 |
|  |  |  | T | 36 | 1.281 | 0.185 | 0.031 | 0.145 |
|  |  | Offline | F | 36 | 1.314 | 0.200 | 0.033 | 0.153 |
|  |  |  | T | 36 | 1.301 | 0.191 | 0.032 | 0.147 |
| Anodal | Object | Online | F | 36 | 1.038 | 0.195 | 0.033 | 0.188 |
|  |  |  | T | 36 | 1.004 | 0.142 | 0.024 | 0.141 |
|  |  | Offline | F | 36 | 1.007 | 0.170 | 0.028 | 0.169 |
|  |  |  | T | 36 | 0.980 | 0.158 | 0.026 | 0.161 |
|  | Action | Online | F | 36 | 1.301 | 0.197 | 0.033 | 0.151 |
|  |  |  | T | 36 | 1.261 | 0.199 | 0.033 | 0.158 |
|  |  | Offline | F | 36 | 1.268 | 0.176 | 0.029 | 0.139 |
|  |  |  | T | 36 | 1.309 | 0.206 | 0.034 | 0.157 |
