## Appendix 3 for "Regionally specific picture naming benefits of focal tDCS are dependent on baseline performance in older adults"

**Appendix 3. Effects of baseline naming and fluid intelligence on stimulation effects in young adults**

| **Cases** | **Sum of Squares** | **df** | **Mean Square** | **F** | **p** | **η²p** |
| --- | --- | --- | --- | --- | --- | --- |
| Stimulation Type | 0.001 | 1 | 0.001 | 0.047 | 0.828 | < .001 |
| Stimulation Type ✻ Baseline Naming Speed | 0.015 | 1 | 0.015 | 0.513 | 0.476 | 0.008 |
| Stimulation Type ✻ Fluid Intelligence | 0.002 | 1 | 0.002 | 0.054 | 0.818 | < .001 |
| Stimulation Type ✻ Region | 0.004 | 1 | 0.004 | 0.120 | 0.730 | 0.002 |
| Stimulation Type ✻ Region ✻ Fluid Intelligence | < .001 | 1 | < .001 | 0.023 | 0.880 | < .001 |
| Stimulation Type ✻ Region ✻ Baseline Naming Speed | 0.016 | 1 | 0.016 | 0.561 | 0.456 | 0.008 |
| Residuals | 1.938 | 66 | 0.029 |  |  |  |
| Naming Type | 0.079 | 1 | 0.079 | 4.750 | 0.033 | 0.067 |
| Naming Type ✻ Baseline Naming Speed | 0.045 | 1 | 0.045 | 2.684 | 0.106 | 0.039 |
| Naming Type ✻ Fluid Intelligence | 0.008 | 1 | 0.008 | 0.475 | 0.493 | 0.007 |
| Naming Type ✻ Region | 0.012 | 1 | 0.012 | 0.709 | 0.403 | 0.011 |
| Naming Type ✻ Region ✻ Fluid Intelligence | 0.011 | 1 | 0.011 | 0.632 | 0.429 | 0.009 |
| Naming Type ✻ Region ✻ Baseline Naming Speed | 0.003 | 1 | 0.003 | 0.174 | 0.678 | < .001 |
| Residuals | 1.097 | 66 | 0.017 |  |  |  |
| Stimulation Time | 0.037 | 1 | 0.037 | 4.699 | 0.034 | 0.066 |
| Stimulation Time ✻ Baseline Naming Speed | 0.046 | 1 | 0.046 | 5.827 | 0.019 | 0.081 |
| Stimulation Time ✻ Fluid Intelligence | 0.006 | 1 | 0.006 | 0.783 | 0.379 | 0.012 |
| Stimulation Time ✻ Region | 0.070 | 1 | 0.070 | 8.818 | 0.004 | 0.118 |
| Stimulation Time ✻ Region ✻ Fluid Intelligence | 0.093 | 1 | 0.093 | 11.717 | 0.001 | 0.151 |
| Stimulation Time ✻ Region ✻ Baseline Naming Speed | 0.012 | 1 | 0.012 | 1.502 | 0.225 | 0.022 |
| Residuals | 0.523 | 66 | 0.008 |  |  |  |
| Stimulation Type ✻ Naming Type | 0.006 | 1 | 0.006 | 0.553 | 0.460 | 0.008 |
| Stimulation Type ✻ Naming Type ✻ Baseline Naming Speed | < .001 | 1 | < .001 | 0.014 | 0.905 | < .001 |
| Stimulation Type ✻ Naming Type ✻ Fluid Intelligence | 0.014 | 1 | 0.014 | 1.274 | 0.263 | 0.019 |
| Stimulation Type ✻ Naming Type ✻ Region | < .001 | 1 | < .001 | 0.013 | 0.909 | < .001 |
| Stimulation Type ✻ Naming Type ✻ Region ✻ Fluid Intelligence | < .001 | 1 | < .001 | 0.024 | 0.877 | < .001 |
| Stimulation Type ✻ Naming Type ✻ Region ✻ Baseline Naming Speed | < .001 | 1 | < .001 | 0.050 | 0.824 | < .001 |
| Residuals | 0.704 | 66 | 0.011 |  |  |  |
| Stimulation Type ✻ Stimulation Time | 0.008 | 1 | 0.008 | 0.956 | 0.332 | 0.014 |
| Stimulation Type ✻ Stimulation Time ✻ Baseline Naming Speed | 0.017 | 1 | 0.017 | 2.075 | 0.154 | 0.030 |
| Stimulation Type ✻ Stimulation Time ✻ Fluid Intelligence | 0.001 | 1 | 0.001 | 0.178 | 0.674 | 0.003 |
| Stimulation Type ✻ Stimulation Time ✻ Region | 0.015 | 1 | 0.015 | 1.846 | 0.179 | 0.027 |
| Stimulation Type ✻ Stimulation Time ✻ Region ✻ Fluid Intelligence | 0.012 | 1 | 0.012 | 1.464 | 0.231 | 0.022 |
| Stimulation Type ✻ Stimulation Time ✻ Region ✻ Baseline Naming Speed | 0.011 | 1 | 0.011 | 1.399 | 0.241 | 0.021 |
| Residuals | 0.531 | 66 | 0.008 |  |  |  |
| Naming Type ✻ Stimulation Time | 0.002 | 1 | 0.002 | 0.294 | 0.589 | 0.004 |
| Naming Type ✻ Stimulation Time ✻ Baseline Naming Speed | 0.003 | 1 | 0.003 | 0.553 | 0.460 | 0.008 |
| Naming Type ✻ Stimulation Time ✻ Fluid Intelligence | 0.018 | 1 | 0.018 | 3.047 | 0.086 | 0.044 |
| Naming Type ✻ Stimulation Time ✻ Region | < .001 | 1 | < .001 | 0.070 | 0.792 | < .001 |
| Naming Type ✻ Stimulation Time ✻ Region ✻ Fluid Intelligence | < .001 | 1 | < .001 | 0.082 | 0.776 | < .001 |
| Naming Type ✻ Stimulation Time ✻ Region ✻ Baseline Naming Speed | 0.001 | 1 | 0.001 | 0.203 | 0.654 | 0.003 |
| Residuals | 0.379 | 66 | 0.006 |  |  |  |
| Stimulation Type ✻ Naming Type ✻ Stimulation Time | 0.004 | 1 | 0.004 | 0.395 | 0.532 | 0.006 |
| Stimulation Type ✻ Naming Type ✻ Stimulation Time ✻ Baseline Naming Speed | 0.002 | 1 | 0.002 | 0.228 | 0.635 | 0.003 |
| Stimulation Type ✻ Naming Type ✻ Stimulation Time ✻ Fluid Intelligence | 0.004 | 1 | 0.004 | 0.399 | 0.530 | 0.006 |
| Stimulation Type ✻ Naming Type ✻ Stimulation Time ✻ Region | 0.028 | 1 | 0.028 | 2.860 | 0.096 | 0.042 |
| Stimulation Type ✻ Naming Type ✻ Stimulation Time ✻ Region ✻ Fluid Intelligence | 0.015 | 1 | 0.015 | 1.510 | 0.224 | 0.022 |
| Stimulation Type ✻ Naming Type ✻ Stimulation Time ✻ Region ✻ Baseline Naming Speed | 0.026 | 1 | 0.026 | 2.619 | 0.110 | 0.038 |
| Residuals | 0.648 | 66 | 0.010 |  |  |  |

*Note.*  Type III Sum of Square

**Between Subjects Effects**

| **Cases** | **Sum of Squares** | **df** | **Mean Square** | **F** | **p** | **η²p** |
| --- | --- | --- | --- | --- | --- | --- |
| Baseline Naming Speed | 6.197 | 1 | 6.197 | 46.441 | < .001 | 0.413 |
| Fluid Intelligence | 0.251 | 1 | 0.251 | 1.882 | 0.175 | 0.028 |
| Region | 0.011 | 1 | 0.011 | 0.083 | 0.774 | 0.001 |
| Region ✻ Fluid Intelligence | 0.153 | 1 | 0.153 | 1.145 | 0.288 | 0.017 |
| Region ✻ Baseline Naming Speed | 0.052 | 1 | 0.052 | 0.391 | 0.534 | 0.006 |
| Residuals | 8.807 | 66 | 0.133 |  |  |  |

*Note.*  Type III Sum of Square
